# Scaling Properties for Artificial Neural Network Models of a Small Nervous System

**DOI:** 10.1101/2024.02.13.580186

**Authors:** Quilee Simeon, Leandro Venâncio, Michael A. Skuhersky, Aran Nayebi, Edward S. Boyden, Guangyu Robert Yang

## Abstract

The nematode worm *C. elegans* provides a unique opportunity for exploring *in silico* data-driven models of a whole nervous system, given its transparency and well-characterized nervous system facilitating a wealth of measurement data from wet-lab experiments. This study explores the scaling properties that may govern learning the underlying neural dynamics of this small nervous system by using artificial neural network (ANN) models. We investigate the accuracy of self-supervised next time-step neural activity prediction as a function of data and models. For data scaling, we report a monotonic log-linear reduction in mean-squared error (MSE) as a function of the amount of neural activity data. For model scaling, we find MSE to be a nonlinear function of the size of the ANN models. Furthermore, we observe that the dataset and model size scaling properties are influenced by the particular choice of model architecture but not by the precise experimental source of the *C. elegans* neural data. Our results fall short of producing long-horizon predictive and generative models of *C. elegans* whole nervous system dynamics but suggest directions to achieve those. In particular our data scaling properties extrapolate that recording more neural activity data is a fruitful near-term approach to obtaining better predictive ANN models of a small nervous system.

## 1. Introduction

Exploring neural system dynamics is crucial in neuroscience and artificial intelligence (AI). This intersection has spurred the evolution of artificial neural network (ANN) models, inspired by biological neural systems. ANNs offer the potential to emulate diverse animal behaviors, providing advantages like detailed specification, causal manipulability, and increasing analytical accessibility, reflecting key aspects of biological nervous systems ([1], [2]). The nematode *Caenorhabditis elegans* (*C. elegans*) is an exemplary model in this context, offering a valuable platform for comparing real and artificial neural dynamics.

*C. elegans* is an excellent model organism for neural dynamics research due to its well-mapped connectome and capabilities for non-invasive neuronal activity tracking via advanced imaging techniques ([3], [4]). The organism’s compact size, transparency, and well-annotated genome allow for intricate optical measurements and deep insights into neural activity. NeuroPAL, a multicolor atlas, allows precise *in vivo* neuron identification, enhancing the capabilities for measurement and analysis of the *C. elegans* nervous system [5].

We formulate the problem of *in silico* nervous system modeling as a teacher-student framework. A real biological neural network (that of *C. elegans*, in our case) is the teacher, ANN models are the students, and self-supervised next time-step neural activity prediction is the curriculum. We consider different instances of the nervous system to be phenotypically matched animals (in our case, adult hermaphrodite worms).

Predicting future neural activity based on historical neural data is not new but the machine learning approach to do it has seen increasing adaption ever since advancements in models like LSTMs demonstrating some success in mammals [6]. In *C. elegans*, the simplified behavioral repertoire and consistent biology offer a unique setting for in-depth model analysis. Self-supervised learning, predicting future states from intrinsic neural patterns, reduces dependence on behaviorally annotated data. While acknowledging the importance of behavior in neural dynamics, our study concentrates on the inherent predictability within neural activity, exploring how neural dynamics can be predicted without direct behavioral reference, similar to how large language models (LLMs) uncover intricate structures in language data [7].

Research into ANNs’ scaling properties has shown that improvements in model size, data volume, and computational resources significantly enhance performance ([8], [9]). The relationship between data size and model capacity is critical in optimizing model performance. However, this relationship in the context of predicting neural dynamics in biological organisms like *C. elegans* is not well-explored. Our study aims to fill this gap by analyzing the impact of data volume, model architecture, and size on ANN performance in neural activity prediction in *C. elegans*. These insights are crucial for optimizing experimental and modeling strategies in neuroscience, contributing to the development of more accurate predictive models for biological nervous systems.

## II. Methods

### A. Neural Activity Data

#### Data sources

We obtained 8 open-source datasets ([5], [10]–[16]) measuring neural activity in *C. elegans* (Table I). These datasets, each recorded under varying experimental conditions, quantify neural activity through the measurement of changes in calcium fluorescence (Δ*f*/*f*_0_) within subsets of the worm’s 302 neurons. The 8 experimental datasets contained different numbers of recorded instances of the nervous system (i.e. worms) as well as variable lengths of time for neural activity measurements (Fig. 1A). The experimental conditions were variable across datasets, ranging from freely moving [10], immobilized [12], and asleep [15] states, to optogenetically stimulated scenarios [11]. However, our modeling strategy is agnostic to the potentially differing conditions and protocols under which the data was acquired as long as the measured system (*C. elegans*) and metric (Δ*F*/*F*_0_) is consistent.

**TABLE 1.**
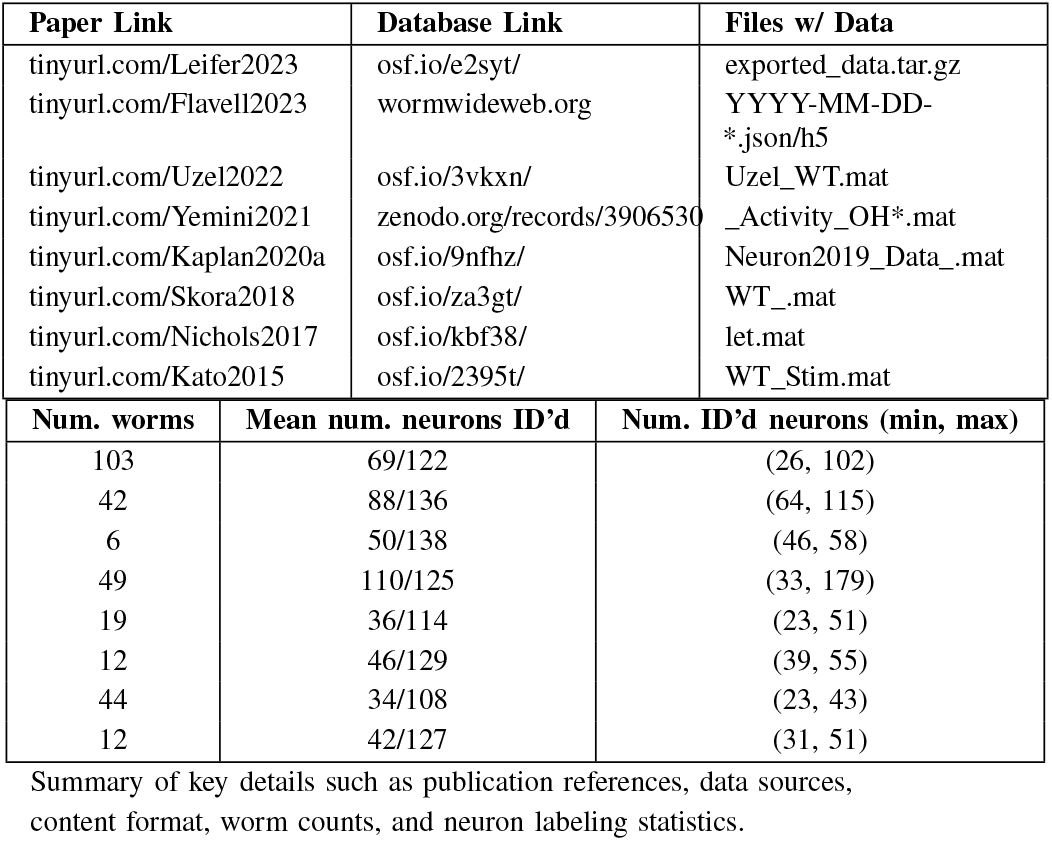
Open-source *C*. *elegans* neural activity datasets.

**Fig. 1.**
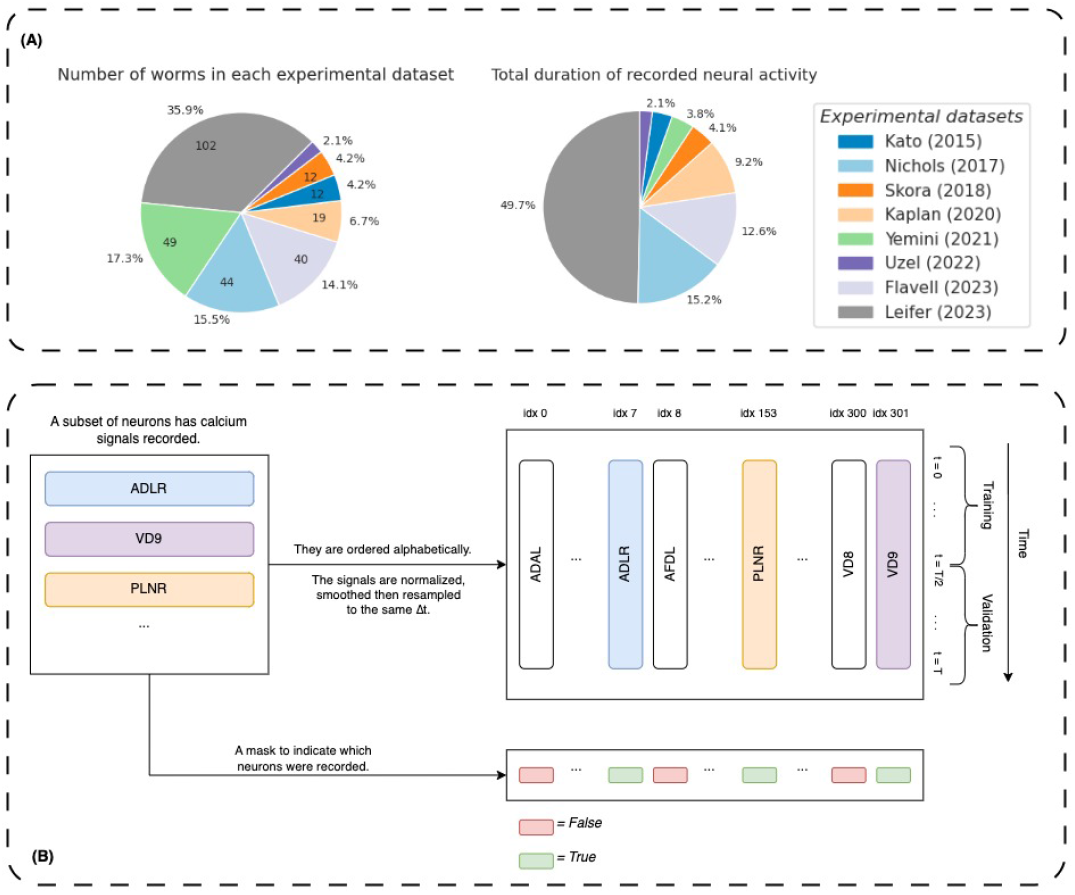
Worm neural datasets curation. Eight open-source *C. elegans* neural activity datasets were downloaded, preprocessed and assimilated. (A) The distribution of the number of worms and the total recording time in each dataset. (B) The neural activity data of all worms is organized a standard format involving a muti-dimensional time series and a boolean feature mask.

#### Standard data format

Each dataset 𝒟^*n*^ includes individual recordings from *n* worms, each consisting of neural activity and a mask indicating the subset of the 302 neurons that were measured and labelled. This mask ensures models are trained only on neural activity recorded from NeuroPAL labelled neurons (Fig. 1B). Specifically:

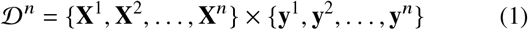

where *n* = |𝒟^*n*^ |.

Each worm, indexed by *k*, has a data matrix **X**^*k*^ ∈ ℝ^302×*T*^*k* and a binary vector **y**^*k*^ ∈{0, 1} ^302^ specifying which neurons were recorded and labelled. Each row 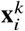 of the matrix **X**^*k*^ contains the time series of neural activity for the *i*^*th*^ neuron for *T*_*k*_ time steps. The rows of **X**^*k*^ and are ordered according to the alphabetical canonical names of the neurons 1 with rows ordered analogously to **y**^*k*^ .

#### Preprocessing

The data, denoted as **X**^*k*^, is processed from the original raw data. First, we normalized the calcium data of each worm by *z*-scoring the full time series independently for each neuron. We then smoothed the signal using a causal exponential kernel (smoothing parameter *α* = 0.5). Finally, the neural data was resampled to a fixed time step interval (Δ*t* ≈ 0.667*s*). Fig. 2 steps through the preprocessing pipeline for a handful of neurons from one worm in the Kato dataset 𝒟_*Kato*_ [16].

**Fig. 2.**
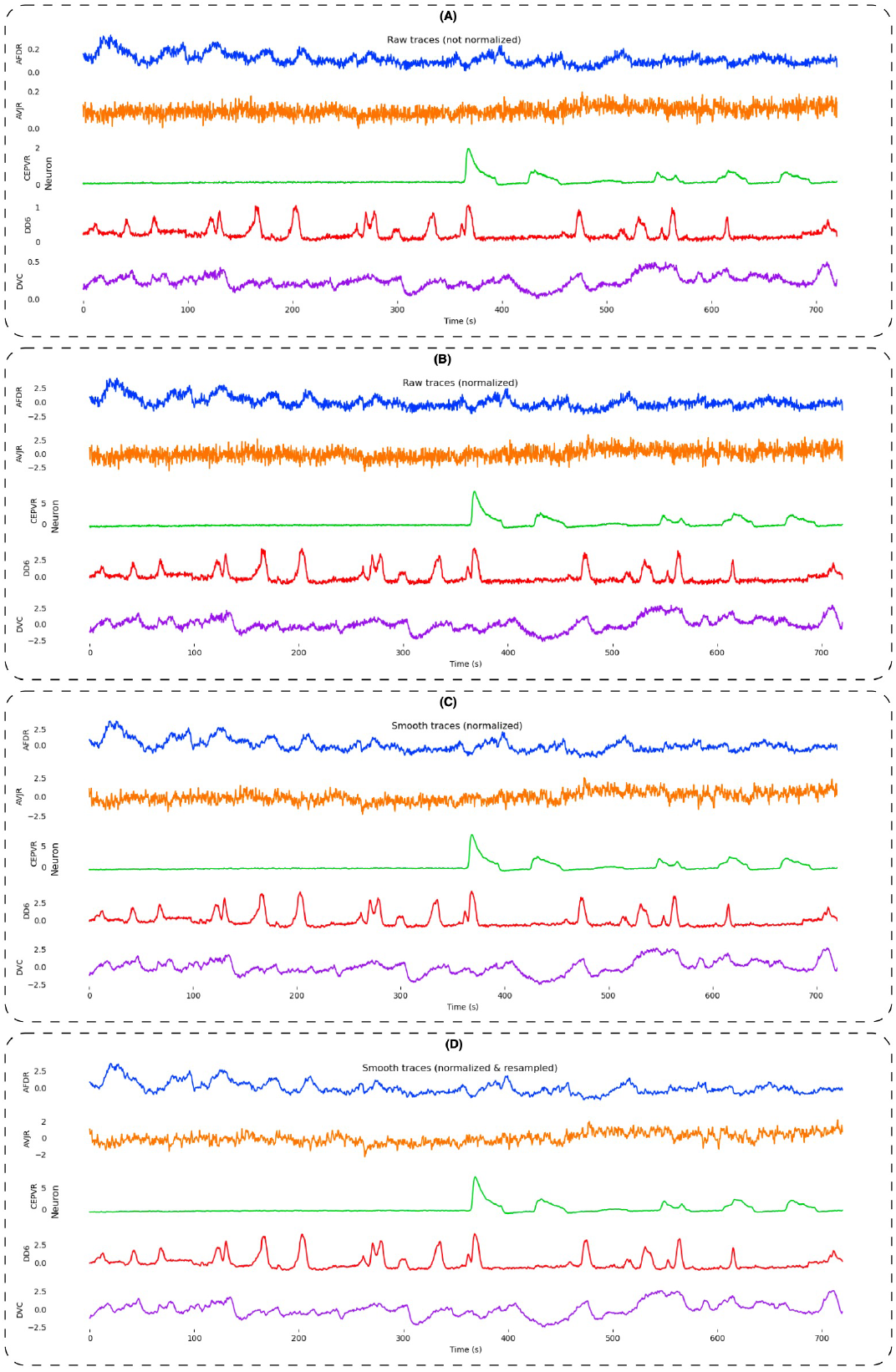
Example of preprocessing worm neural data. We use the example of the Kato (2015) dataset to illustrate steps of the preprocessing the neural activity data. Starting with the raw calcium fluorescence signals (A); we first standardize or *z*-score each neuron independently (B); then smooth using a causal filter (C); and finally resample to a fixed time step (D).

#### Train-Test split

For each worm’s neural activity data matrix **X**^*k*^, we performed a temporal split to create a training set 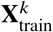 and a testing set 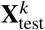 . A balanced 50:50 split was adopted, allocating the first half of the neural activity recording to the training set and the second half to the testing set. One might create equally sized train and test sets containing *n*_s_ sequences of length *L* by sampling their start indices uniformly from (or equidistantly within) the range [0, ⌊*T*/2⌋ − *L* − 1) and [⌈*T*/2⌉, *T* − *L* − 1] for train and test, respectively

#### Amount of Data

The ability to vary the amount of training data is central to our investigation of the effects of data scaling on the ability of self-supervised models to do future neural activity prediction. However, we are data constrained in our setting since the collective dataset of all worms 𝒟_*ALL*_ has only 284 worms (Fig. 3).

**Fig. 3.**
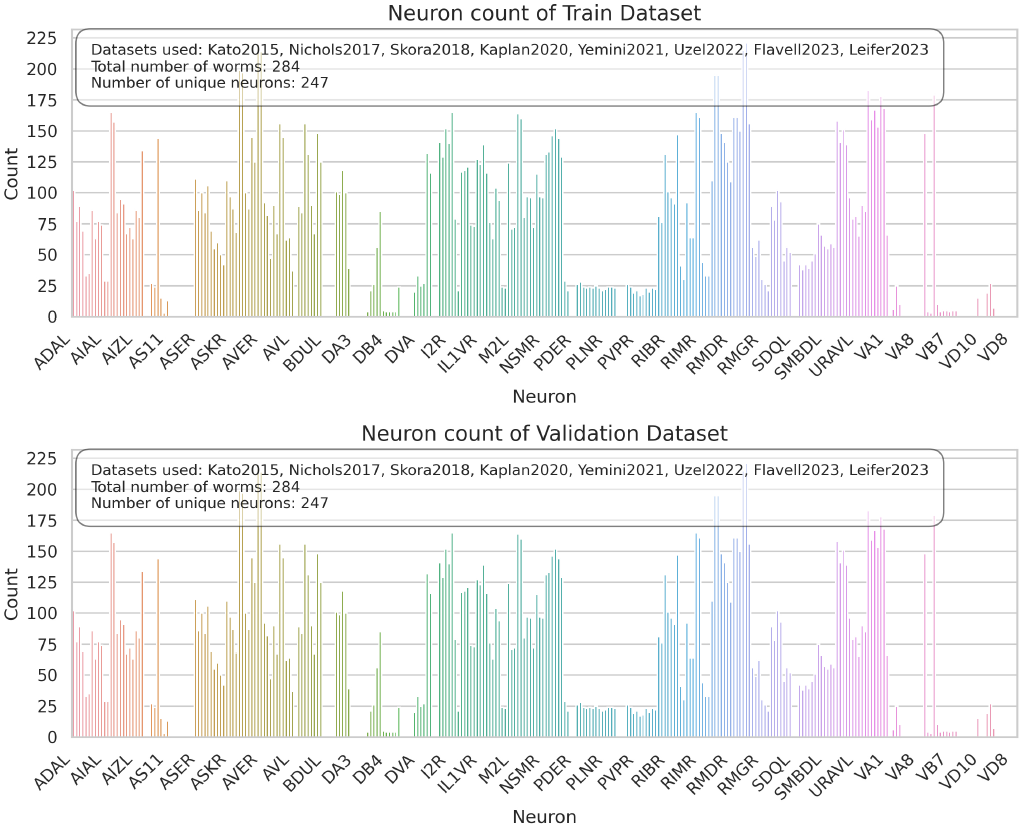
Distribution of neurons in 𝒟_*ALL*_. Since the train-validation split is along the temporal (not feature) dimension, the neuron distribution is the same in the train and validation sets. The combined dataset has 284 worms from the 8 experimental dataset sources, with 247 out the 302 neuron classes in *C*.*elegans* represented (i.e. recorded in at least 1 worm). Of the recorded neurons, some are over-represented whereas others have been recorded in only 1 worm.

#### Source Datasets

The experimental sources contributing to 𝒟_*ALL*_ = 𝒟^284^ are Kato 𝒟_*Kato*_ (|𝒟_*Kato*_| = 12), Nichols 𝒟_*Nichols*_ (|𝒟_*Nichols*_| = 44), Skora 𝒟_*Skora*_ (|𝒟_*Skora*_| = 12), Kaplan 𝒟_*Kaplan*_ (|𝒟_*Kaplan*_| = 19), Yemini 𝒟_*Yemini*_ (|𝒟_*Yemini*_| = 49), Uzel 𝒟_*Uzel*_ (|𝒟_*Uzel*_| = 6), Flavell 𝒟_*Flavell*_ (|𝒟_*Flavell*_| = 42), Leifer 𝒟_*Leifer*_ (|𝒟_*Leifer*_| = 103).

#### Mixed Datasets

To create mixed datasets combining worms from the different experimental sources, we randomly sample from the combined pool of all available worms 𝒟_*ALL*_.

Let *i* ∈ [8] := {1, 2, …, 8} index into the list of experimental sources (sorted by publication date): [Kato, Nichols, …, Leifer]. We denote a mixed worm dataset containing *n*_*w*_ worms from any combination of experimental sources as 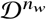.

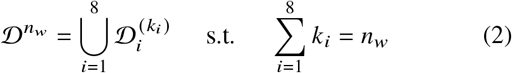

Here, 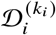 stands for *k*_*i*_ worms sampled specifically from the experimental dataset indexed by *i* (Fig. 4A). Note that there are multiple assignments that can achieve a dataset with *n*_*w*_ worms. Therefore, 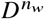 is a random variable. Our mixed dataset sampling process is akin to sampling from a multinomial distribution where the probabilities correspond to the proportion of available worms from each experimental source:

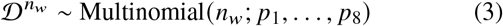

where *p*_*i*_ reflects the relative contribution of the experimental dataset indexed by *i* to the pool.

**Fig. 4.**
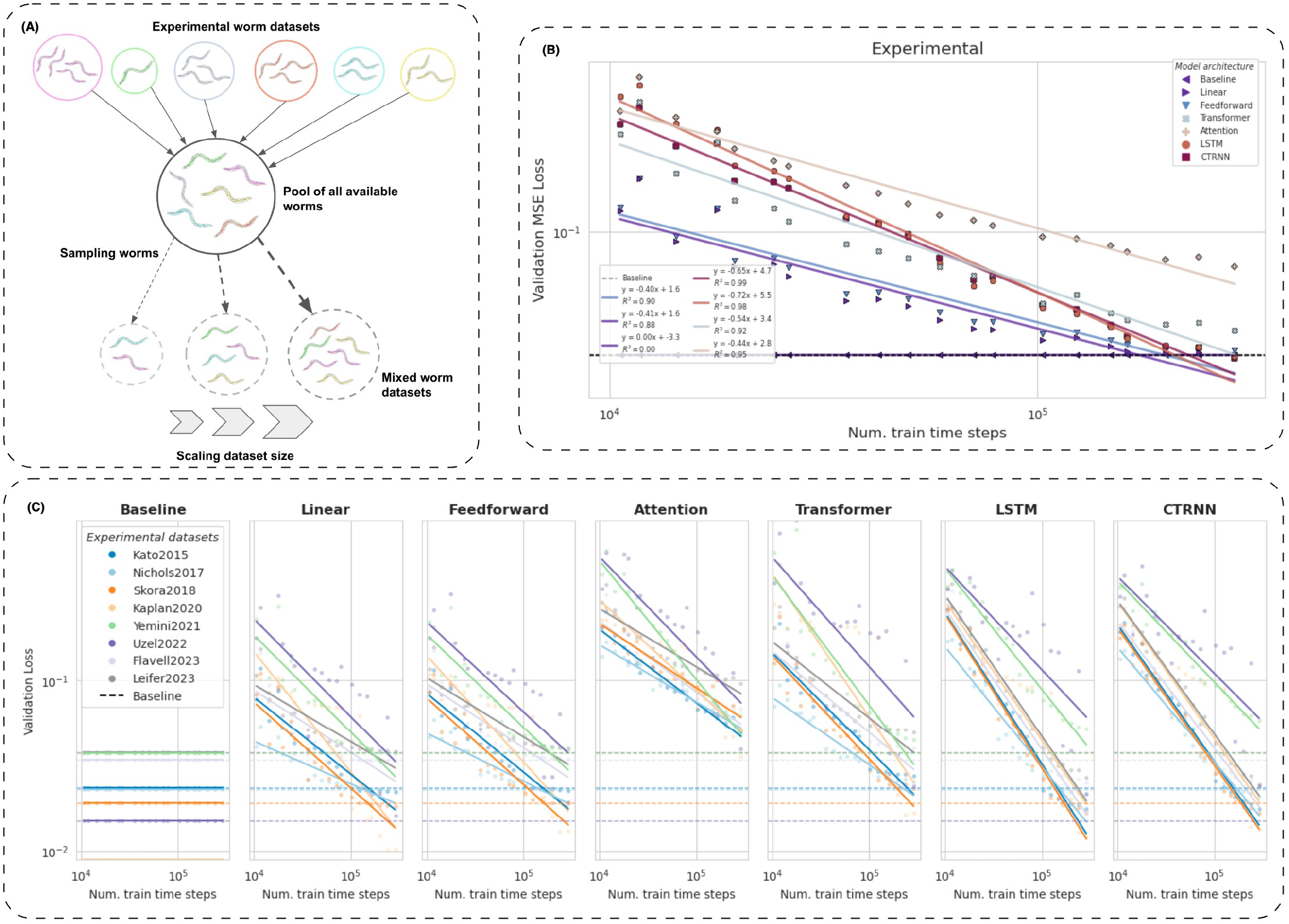
Sampling datasets and dataset scaling experiment. (A) Creating mixed worm datasets of various sizes by sampling from the pool of all available worms across different experimental datasets. (B) The validation loss from optimizing for next time step prediction is a decreasing function of the number of training time steps and depends on the model architecture. All models are approximately matched in size at 580K (K=thousand) trainable parameters. (C) Within a model class, the data scaling slopes exhibit remarkable similarity across experimental dataset sources, despite the diversity of the experimental and behavioral conditions.

This methodical approach allows us to create increasingly larger mixed datasets up to the largest one containing all worms 𝒟_*ALL*_ := 𝒟^284^ (for which there is only 1 possible assignment). The result is a series of mixed datasets, each with a unique composition of worms, yet collectively spanning the full range of neural dynamics present in the collective data. The mixed datasets 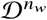 thus represent a diverse cross-section of neural activities encompassing variations in experimental conditions.

We could also generate subsets from any single experimental dataset using this approach by simply restricting our random sampling to that source. This allows us to create increasingly larger data subsets constrained to a particular experimental context.

#### Synthetic Datasets

We created synthetic datasets that mirror the complexity and challenges of the processed real *C*.*elegans* neural activity data, such as partial observability, sparsity, and noise, with 200 synthetic ‘worms’ each having 1500 time steps of activity from 50 randomly chosen neurons. These datasets allow for robust model validation against known dynamics, offering clear benchmarks for performance evaluation.

The **Sines** dataset models each neuron’s activity as an independent sinusoid with a random phase but consistent frequency across worms, exploring an assumption of neuron-specific ‘fingerprints’ in the network dynamics. This dataset tests models on their ability to capture simple, uncoupled oscillatory patterns, a task suited for recurrent models like LSTMs.

Conversely, the **Random Walk** dataset challenges models with inherently unpredictable neural activity simulated through a random walk process, aiming to benchmark against the naive predictor’s performance which is provably the best estimator in this case. This setup serves as a critical test of model implementations, ensuring no model unjustifiably surpasses this baseline.

### B. Model Structure

#### Model architectures

Our study utilizes three distinct classes of neural networks to harness different inductive biases for the prediction of future neural activity in *C. elegans*. These include Long-Short Term Memory (LSTM) networks, Transformer networks, and Feedforward networks. These architectures were chosen to represent a fundamental set of mechanisms—recurrence, attention, and feedforward processing—allowing us to assess the impact of structural and mechanistic differences on the task at hand.

#### Shared model structure

Each architecture is implemented within a common structural framework comprising an embedding block, a hidden ‘core’ module, and a linear readout layer to enable a consistent training and evaluation procedure.

This shared structure is modular allowing for the comparison of different ANN architectures by substituting only the core module (Fig. 5A).

**Fig. 5.**
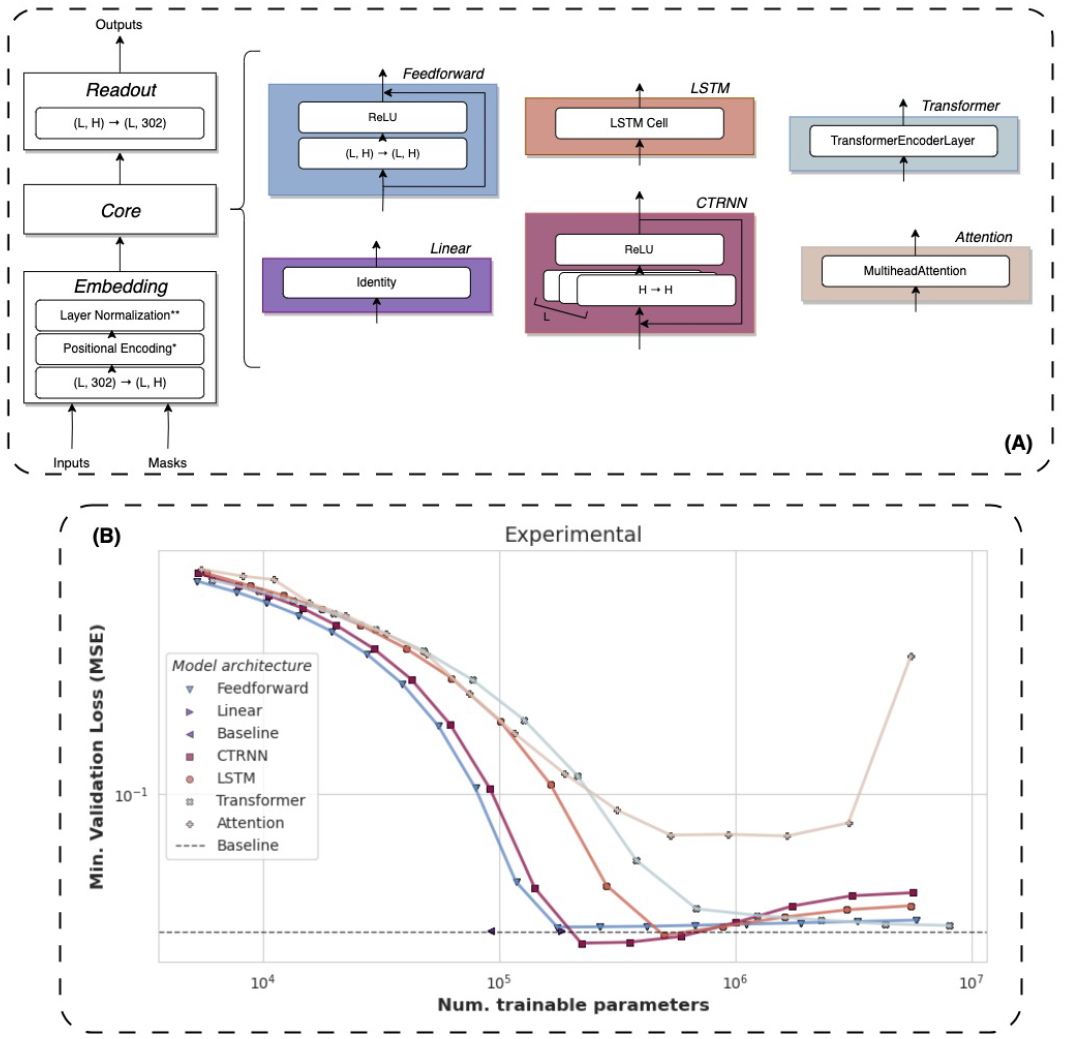
Model architecture and model scaling experiment. (A) The six (6) model classes we investigate share a common backbone but have differentiated core architectures: Feedforward, Linear, LSTM, CTRNN, Transformer, and Attention. Asterisks indicate modules appear in some architectures but not others. *Positional encoding present here in the Transformer and Attention models. **Layer normalization absent here in the Feedforward and Attention models. (B) The next time step prediction validation accuracy (inverse relation to loss) improves non-linearly with increasing the model sizes with some model architectures scaling better than others. Only the recurrent (CTRNN and LSTM) models beat the baseline loss. All models are trained and validated on the fixed largest training and validation datasets, respectively, made from 𝒟_*ALL*_.

1. *Embedding:* The basic embedding layer linearly projects from the 302-dimensional neural state space to a higher or lower *H*-dimensional latent space. Certain architectures like the Transformer may add a positional encoding and other architectures may apply layer normalization to stabilize the learning process.
2. *Core:* The core module is architecture-specific and constitutes the primary computational engine of the model. It is restricted to use a single layer to maintain simplicity and facilitate interpretability. Besides the single layer restriction, we place no constraints on the class of computations the core is allowed to use (e.g. recurrence, parallel-processing, etc.), making our approach highly modular and scalable. The Baseline and LinearRegression prediction models are ‘shallow’ models, which means that they lack the ‘core’ module.
3. *Output Mapping:* The final component of the model is a linear projection from the latent space back to the original neural state space. The output of this layer is the predicted future neural activity 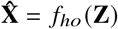, where *f*_*ho*_ is the linear transformation from the hidden to the output space.

#### Causal Predictions and Temporal Memory

Our models are tasked with making causal predictions, where future predictions do not rely on future inputs. We use a causal attention mask in the Transformer and Attention models. The LSTM and CTRNN models are inherently causal by definition. The Linear and Feedforward models lack access to temporal context beyond the current time step (i.e. they process each time point in a sequence independently). This essentially restricts the computation of the Linear and Feedforward models to feature regression (linear and nonlinear, respectively), providing a baseline for the importance of temporal information in self-supervised neural prediction.

**Baseline Model**

We use the naive predictor which posits that the next neural state will be identical to the current one as our baseline model. This baseline is a commonly used one for time series prediction tasks and it is the known optimal predictor for a random walk. Despite its simplicity, this baseline is not trivial to beat in the context of neural activity data which can often resemble, at a first-order approximation, a random process. Beating this baseline requires our ANN models to uncover and leverage complex, higher-order structures in the neural activity data beyond what is expected from a purely stochastic process. To maintain consistency with our models super-structure (Fig. 5A left), our Baseline model is implemented as’thunk’ model class with no trainable parameters that simply copies its masked input as its output.

### C. Training Objective and Loss Function

#### Training Objective

The models are trained under a selfsupervised objective to predict the 1-time step shifted sequence of neural activity given a input sequence of neural activity of length *L*. The training objective is simply to minimize the mean-squared error (MSE) between the predicted and true neural activity sequences. The loss function further incorporates the boolean neuron mask to ensure that only neurons with measured data contribute to the loss computation. The mean-squared error (MSE) loss function with the boolean mask is defined as:

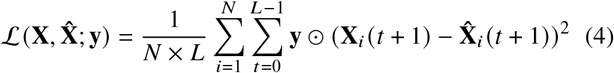

where **X**_***i***_ (*t*+1) is the true activity of the *i*^*th*^ neuron at time 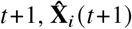 the predicted activity at time *t* + 1, **y** ∈ {0, 1} ^302^ is the boolean feature mask indicating the presence of data for neuron *i, L* is the sequence length used for training the model, and *N* = 𝟙^⊤^**y** is the number of masked neurons (i.e. the number of labelled neurons with data).

#### Data Sampling and Model Evaluation

We construct the training and validation sets by sampling from each worm *n*_*s*_ = 32 sequences of length *L* = 180 time steps according to the method in subsection II-A **Train-Test split**. For example, a dataset 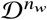 containing *n*_*w*_ worms would produce training and validation sets each with *n*_*w*_ × 32 sequences. Train and validation data loaders use a batch size of 128.

#### Training Protocol

Models are trained up to a maximum of 500 epochs using the AdamW optimizer, with an initial learning rate of 0.001. A learning rate scheduler reduces the rate upon a validation loss plateau, with a decay factor of 0.1. Early stopping with a patience of 100 epochs is employed for efficiency. Training for all experiments was run with the same computing resources and device specifications (1 NVIDIA A100 80GB GPU).

## III. Results

### A. Data Scaling

#### 1). Mixed Dataset Scaling

To assess how increasing the amount of training data influences the self-supervised next time step neural activity prediction.

We trained models of every architecture on incrementally larger training sets from a sequence of mixed worm datasets ranging in size from 1 to 284 worms, sampled according to subsection III-A1 **Mixed Dataset Scaling**. At each training set size, the models were evaluated against the fixed maximum sized validation set made from 𝒟_*ALL*_. Since there are multiple possible combinatorial assignments for a mixed dataset 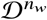 of size *n*_*w*_ < 284 worms, we plot the independent variable as the number time-steps in the train dataset. We controlled for the hidden size of each model architecture class so that all models were approximately matched at 580K (K=thousand) trainable parameters (Table II).

**TABLE 2.**
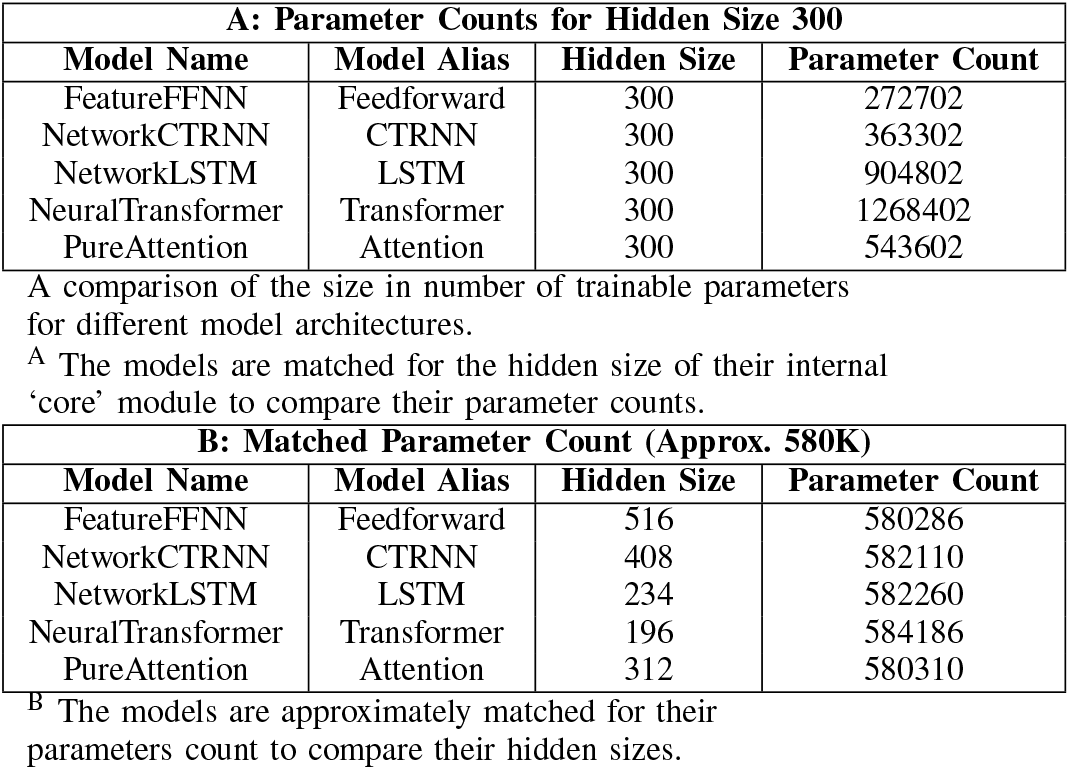
Model Hidden Sizes and Parameter Counts.

The results shown in Fig 4B indicate that, at a fixed model size 0.580M trainable parameters, the CTRNN models scale to scale the best (slope = − 0.65) with dataset size, whereas the Feedforward models scale the worst (slope = − 0.40), out of the architecture classes investigated. We also validated our dataset scaling on the synthetic datasets introduced in subsection II-A **Synthetic Datasets** (Fig. 6A-B).

**Fig. 6.**
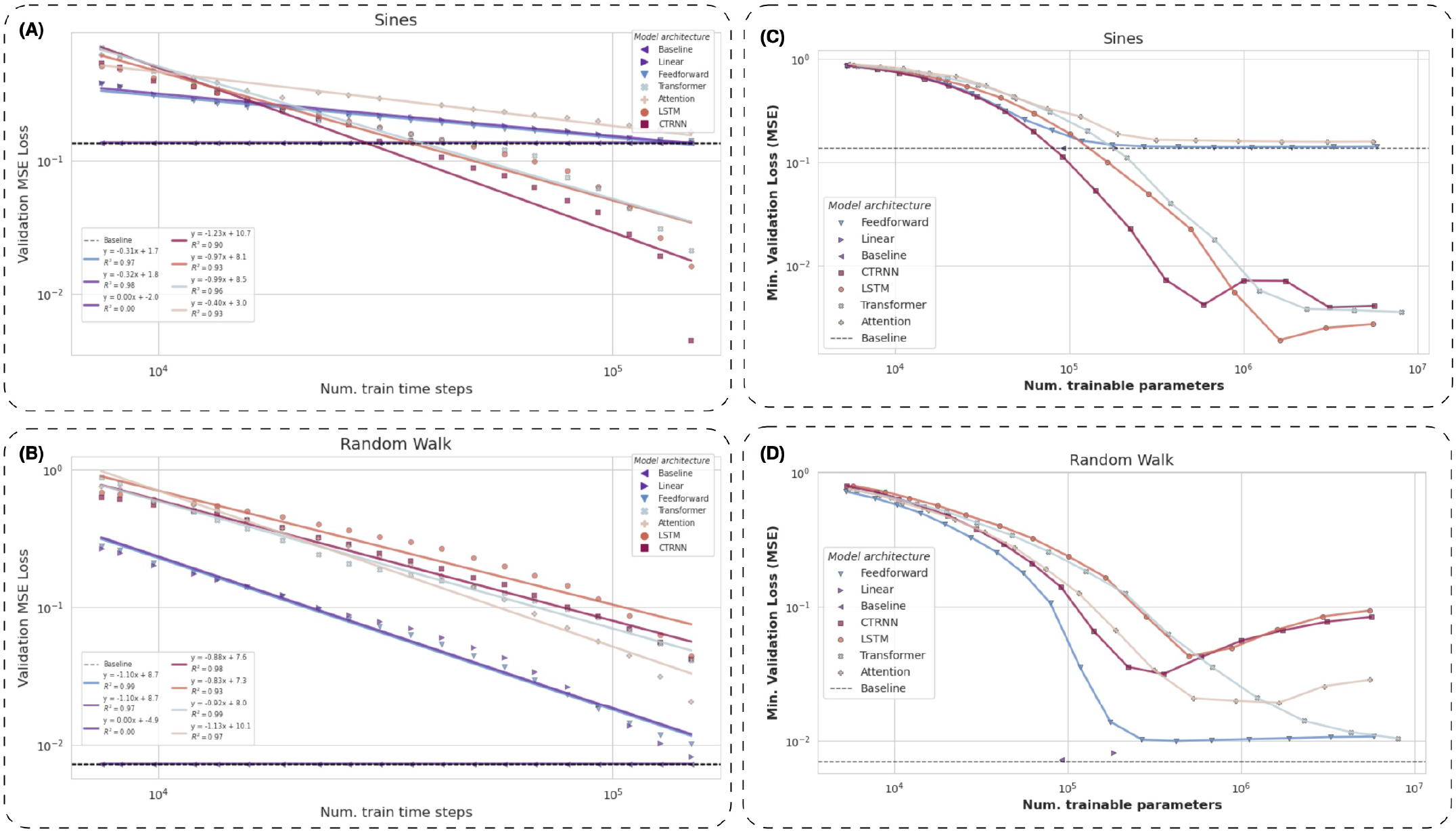
Dataset and model size scaling on synthetic datasets. (A-B) Dataset size scaling experiments. (A) Dataset size scaling of the ANN models used in 4B on the Sines dataset. Beyond a modest dataset size threshold, the sequence models – recurrent and transformer – outperform the naive predictor baseline loss. (B) Dataset size scaling of the ANN models used in 4B on the Random Walk dataset. As expected, no model beats the naive predictor baseline loss since that is the optimal predictor for a random walk. (C-D) Model size scaling experiments. (C) Model size scaling of the ANN models used in 5B on the Sines dataset. Beyond modest model size threshold, the sequence models – recurrent and transformer – outperform the naive predictor baseline loss. (D) Model size scaling of the ANN models used in 5B on the Random Walk dataset. No model can beat the naive predictor baseline loss, even after training on all the available data.

#### 2) Individual Dataset Scaling

To determine if models trained on mixed datasets maintained consistent scaling properties when evaluated on the individual experimental source datasets.

Utilizing the best model from the mixed dataset scaling experiment at each training dataset size, we evaluated on the largest validation set made from each of the experimental sources (refer to last paragraph of subsection II-A **Mixed Datasets**).

Fig. 4C presents the results for scaling the sizes of the individual experimental source datasets.

### B. Model Scaling

To determine the effect of model complexity, as determined by the number of trainable parameters and architecture, on the performance of self-supervised neural activity prediction in *C. elegans*.

We varied the hidden size of the ‘core’ architecture as a knob with which to vary the number of parameters of our models. Since each architecture (Linear, Feedforward, CTRNN, LSTM, Transformer, Attention) has a different number of trainable parameters than the others at any given hidden size, we plot the number of trainable parameters as the independent variable. The various sized models of each architecture/class were trained on the same, fixed maximum sized training dataset made from 𝒟_*ALL*_. We also validated our model scaling on the synthetic datasets introduced in subsection II-A **Synthetic Datasets** (Fig. 6C-D).

Fig. 5B shows the results of increasing the model size for the different architecture classes. As discussed in subsection II-B **Shared model structure**, the Baseline and Linear models have no hidden size dimension to vary. The Baseline model simply has no trainable parameters at all, whereas the Linear model has no trainable parameters in its ‘core’ module – which is just an Identity layer (Fig. 5A left).

The Baseline model achieved a minimum validation loss of 0.03541 which, by definition, is the baseline loss. The Linear model achieved a minimum validation loss of 0.03533. The minimum validation loss and parameter counts of the other trained models is presented in Fig. 7, along with a qualitative comparison of their auto-regressive generation capability.

**Fig. 7.**
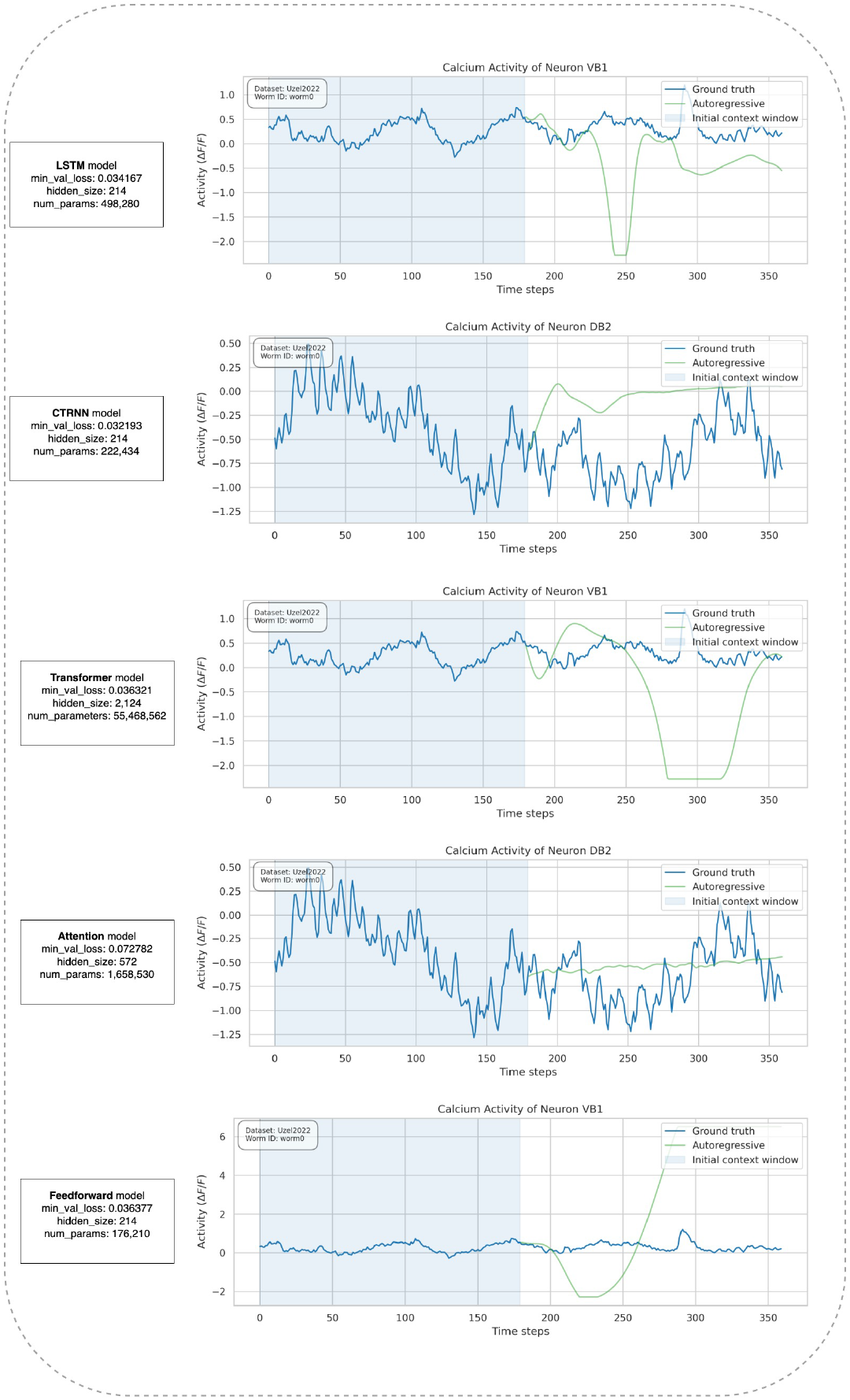
Auto-regressive generation of trained models on 𝒟_*Uzel*_. Models attaining the lowest validation loss for some architecture classes in Fig. 5B are seeded with a length *L* = 180 time step sequence from the validation set of one worm in 𝒟_*Uzel*_ as context. The models are then made to autoregressively generate the next 180 time-steps by repeatedly predicting 1-time step ahead and appending this prediction to the end of a sliding context window.

### C. Synthetic Experiments

To investigate the degree to which the scaling properties observed for the real worm datasets are a function of the underlying system (*C elegans*) versus of the networks being trained.

We trained the same ANN classes and sizes used to model the real *C. elegans* neural datasets (sections III-A and III-B) on two synthetic neural datasets generated by known dynamical systems: a sinusoidal oscillator (Sines) and a stochastic process (Random Walk).

To scale the dataset sizes, for each architecture class, we sampled increasingly larger subsets of worms from each synthetic dataset (Sines or Random Walk) ranging from 1 worm to all 200 worms. Varying the number of worms in a dataset was an indirect way to vary the number of time steps of neural activity in the dataset since those two variables are positively correlated. The number of time steps, in turn, causally determines the amount of data used for training. To scale the model sizes, for each architecture class, we increased the number of parameters by varying the hidden size in the range [8, 4096] at log-spaced intervals. The hidden size was a proxy variable that could be manipulated to directly vary the number of trainable parameters in the models.

For the dataset size scaling experiments, all models were approximately matched in size at 580K (K=thousand) trainable parameters and the validation set used was the fixed largest possible one made from all 200 worms in the synthetic dataset. For the model size scaling experiments, the training and validation sets used were the fixed largest possible ones made from all 200 worms in the synthetic dataset.

Fig.6 presents the results for scaling the sizes of the synthetic datasets.

## IV. Discussion

This study’s exploration into the scaling properties of ANNs in predicting neural activity within *C. elegans* reveals critical insights into data-driven modeling of biological neural networks. Our results demonstrate that the predictive accuracy of ANNs is significantly influenced by both the volume of training data and the complexity of the model. We observed a logarithmic decrease in mean-squared error with increased training data, a consistency that persisted across various experimental datasets. This suggests a pivotal role for data volume in enhancing model performance, with further gains possible through optimal model complexity.

Model architecture emerged as a decisive factor; recurrent models, such as LSTMs and CTRNNs, outperformed others, underscoring the importance of temporal dynamics in neural activity prediction. The experiments indicated that different datasets, despite their diverse origins, share common underlying dynamics, as evidenced by the similar scaling slopes in prediction accuracy. This finding substantiates the pooling of data from multiple sources to improve model robustness.

Challenges encountered include determining appropriate model sizes for varying data volumes and incorporating behavioral contexts into predictions, which present promising avenues for future research. Our work thus far has not fully realized the long-horizon predictive capabilities for *C. elegans* neural dynamics. However, it paves the way for more comprehensive models by highlighting the potential benefits of incorporating richer datasets and nuanced model architectures.

Future efforts will focus on refining models to capture the complexity of neural dynamics more accurately. This includes considering the integration of behaviorally-annotated data to provide additional context and investigating architectures capable of leveraging larger datasets without succumbing to the diminishing returns of over-complexity. Extension of these approaches to more complex nervous systems could offer valuable comparative insights and further the understanding of neural dynamics prediction.

In conclusion, this research contributes foundational knowledge towards the development of ANN models that more accurately reflect the intricacies of biological neural networks, bridging the gap from model organisms to broader biological contexts.

## V. Reproducibility

We have made the combined *C. elegans* neural activity dataset publicly available on the Hugging Face platform here: qsimeon/celegans_neural_data. All the code written for this study has been released publicly on GitHub at this repository: metaconsciousgroup/worm-graph.

## Acknowledgment

We would like to thank our undergraduate interns, Kaiya Ivy Zhao and Tiffney Aina, for their invaluable contributions to early, exploratory work on this project.

## Footnotes

1 https://www.wormatlas.org/NeuronNames.htm

## Notes

https://huggingface.co/datasets/qsimeon/celegans_neural_data

https://github.com/metaconsciousgroup/worm-graph.git

### Competing Interest Statement

The authors have declared no competing interest.

### Summary of Updates

Reformatted in IEEE conference paper style, added figures, removed appendix, reordered authors.

## References

[1] D. L. K. Yamins and J. J. DiCarlo, “Using goal-driven deep learning models to understand sensory cortex,” Nat. Neurosci., vol. 19, no. 3, pp. 356–365, Mar. 2016.

[2] D. L. K. Yamins, H. Hong, C. F. Cadieu, E. A. Solomon, D. Seibert, and J. J. DiCarlo, “Performance-optimized hierarchical models predict neural responses in higher visual cortex,” Proc. Natl. Acad. Sci. U. S. A., vol. 111, no. 23, pp. 8619–8624, Jun. 2014.

[3] A. M. Leifer, C. Fang-Yen, M. Gershow, M. J. Alkema, and A. D. T. Samuel, “Optogenetic manipulation of neural activity in freely moving Caenorhabditis elegans,” Nat. Methods, vol. 8, no. 2, pp. 147–152, Feb. 2011, doi: 10.1038/nmeth.1554.

[4] J. P. Nguyen et al., “Whole-brain calcium imaging with cellular resolution in freely behaving Caenorhabditis elegans,” Proceedings of the National Academy of Sciences, vol. 113, no. 8, pp. E1074–E1081, 2016, doi: 10.1073/pnas.1507110112.

[5] E. Yemini, A. Lin, A. Nejatbakhsh, E. Varol, R. Sun, G. E. Mena, A. D. T. Samuel, L. Paninski, V. Venkatachalam, and O. Hobert, “NeuroPAL: A Multicolor Atlas for Whole-Brain Neuronal Identification in C. elegans,” Cell, vol. 184, pp. 272–288.e11, 2021, doi: 10.1016/j.cell.2020.12.012.

[6] C. Pandarinath et al., “Inferring Single-Trial Neural Population Dynamics Using Sequential Auto-Encoders,” Nature Methods, vol. 15, no. 10, pp. 805–815, 2018.

[7] A. Radford et al., “Language Models are Unsupervised Multitask Learners,” OpenAI Blog, vol. 1, no. 8, 2019.

[8] J. Kaplan et al., “Scaling Laws for Neural Language Models,” arXiv [cs.LG], 2020, 2001.08361.

[9] J. Hoffmann et al., “Training Compute-Optimal Large Language Models,” arXiv [cs.CL], 2022, 2203.15556.

[10] A. A. Atanas et al., “Brain-wide Representations of Behavior Spanning Multiple Timescales and States in C. elegans,” Cell, vol. 186, no. 19, pp. 4134–4151.e31, Sep. 2023.

[11] F. Randi et al., “Neural Signal Propagation Atlas of C. elegans,” arXiv preprint 2208.04790, 2022.

[12] K. Uzel, S. Kato, and M. Zimmer, “A set of hub neurons and non-local connectivity features support global brain dynamics in C. elegans,” Current Biology, vol. 32, no. 16, pp. 3443–3459, 2022, doi: 10.1016/j.cub.2022.06.039.

[13] H. S. Kaplan, O. S. Thula, N. Khoss, and M. Zimmer, “Nested neuronal dynamics orchestrate a behavioral hierarchy across timescales,” Neuron, vol. 105, no. 3, pp. 562–576, 2020.

[14] S. Skora, F. Mende, and M. Zimmer, “Energy scarcity promotes a brain-wide sleep state modulated by insulin signaling in C. elegans,” Cell reports, vol. 22, no. 4, pp. 953–966, 2018.

[15] A. L. A. Nichols et al., “A global brain state underlies c. elegans sleep behavior,” Science, vol. 356, no. 6344, pp. eaam6851, 2017.

[16] S. Kato, H. S. Kaplan, T. Schrödel, S. Skora, T. H. Lindsay, E. Yemini, S. Lockery, and M. Zimmer, “Global brain dynamics embed the motor command sequence of Caenorhabditis elegans,” Cell, vol. 163, no. 3, pp. 656–669, Oct. 2015, doi: 10.1016/j.cell.2015.09.034.

[17] A. S. Abdelfattah et al., “Neurophotonic tools for microscopic measurements and manipulation: status report,” Neurophotonics, vol. 9, p. 013001, 2022.

[18] D. G. Albertson, J. N. Thompson, and S. Brenner, “The Pharynx of Caenorhabditis Elegans,” Philosophical Transactions of the Royal Society of London. Series B, Biological Sciences, vol. 275, no. 938, pp. 299–325, 1997.

[19] N. Agarwal, N. Mehta, A. C. Parker, and K. Ashouri, “C. elegans neuromorphic neural network exhibiting undulating locomotion,” in 2017 International Joint Conference on Neural Networks (ĲCNN), IEEE, 2017.

[20] I. Beets et al., “System-wide mapping of neuropeptide-GPCR interactions in C. elegans,” bioRxiv, 2022.10.30.514428, 2022.

[21] B. Bentley et al., “The Multilayer Connectome of Caenorhabditis elegans,” PLoS Comput. Biol., vol. 12, e1005283, 2016.

[22] J. H. Boyle and N. Cohen, “Caenorhabditis elegans body wall muscles are simple actuators,” Biosystems, vol. 94, pp. 170–181, 2008.

[23] J. H. Boyle, S. Berri, and N. Cohen, “Gait Modulation in C. elegans: An Integrated Neuromechanical Model,” Front. Comput. Neurosci., vol. 6, p. 10, 2012, doi: 10.3389/fncom.2012.00010.

[24] C. A. Brittin, S. J. Cook, D. H. Hall, S. W. Emmons, and N. Cohen, “A multi-scale brain map derived from whole-brain volumetric reconstructions,” Nature, vol. 591, pp. 105–110, 2021, doi: 10.1038/s41586-021-03284-x.

[25] T. B. Brown et al., “Language Models are Few-Shot Learners,” arXiv preprint 2005.14165, 2020.

[26] B. L. Chen, D. H. Hall, and D. B. Chklovskii, “Wiring optimization can relate neuronal structure and function,” Proc. Natl. Acad. Sci. U. S. A., vol. 103, pp. 4723–4728, 2006, doi: 10.1073/pnas.0506806103.

[27] S. J. Cook et al., “Whole-animal connectomes of both Caenorhabditis elegans sexes,” Nature, vol. 571, no. 7763, pp. 63–71, 2019.

[28] U. Dag et al., “Dissecting the functional organization of the C. elegans serotonergic system at whole-brain scale,” Cell, 2023, doi: 10.1016/j.cell.2023.04.023.

[29] G. T. Einevoll et al., “The Scientific Case for Brain Simulations,” Neuron, vol. 102, pp. 735–744, 2019, doi: 10.1016/j.neuron.2019.03.027.

[30] C. Eliasmith and O. Trujillo, “The use and abuse of large-scale brain models,” Curr. Opin. Neurobiol., vol. 25, pp. 1–6, 2014, doi: 10.1016/j.conb.2013.09.009.

[31] K. Funahashi and Y. Nakamura, “Approximation of dynamical systems by continuous time recurrent neural networks,” Neural Netw., vol. 6, no. 6, pp. 801–806, Jan. 1993.

[32] D. H. Hall and Z. F. Altun, C. Elegans Atlas, Cold Spring Harbor, NY, USA: CSHL Press, 2008.

[33] D. Harel, “A grand challenge: Full reactive modeling of a multi-cellular animal,” in Hybrid Systems: Computation and Control Lecture notes in computer science, Springer Berlin Heidelberg, 2003, pp. 2–2, doi: 10.1007/3-540-36580-x_2.

[34] M. Hendricks, H. Ha, N. Maffey, and Y. Zhang, “Compartmentalized calcium dynamics in a C. elegans interneuron encode head movement,” Nature, vol. 487, pp. 99–103, 2012, doi: 10.1038/nature11081.

[35] S. Herculano-Houzel, B. Mota, and R. Lent, “Cellular scaling rules for rodent brains,” Proc. Natl. Acad. Sci. U. S. A., vol. 103, pp. 12138–12143, 2006, doi: 10.1073/pnas.0604911103.

[36] J. Hestness et al., “Deep Learning Scaling Is Predictable, Empirically,” arXiv [cs.LG], 2017, 1712.00409.

[37] G. E. Hinton, “Connectionist learning procedures,” Artificial intelligence, vol. 40, no. 1-3, pp. 185–234, 1989.

[38] O. Hobert, L. Glenwinkel, and J. White, “Revisiting Neuronal Cell Type Classification in Caenorhabditis Elegans,” Current Biology: CB, vol. 26, no. 22, pp. R1197–R1203, 2016, doi: 10.1016/j.cub.2016.10.027.

[39] S. Hochreiter and J. Schmidhuber, “Long Short-Term Memory,” Neural Computation, vol. 9, no. 8, pp. 1735–1780, 1997, doi: 10.1162/neco.1997.9.8.1735.

[40] T. R. Insel, S. C. Landis, and F. S. Collins, “Research priorities. The NIH BRAIN Initiative,” Science, vol. 340, pp. 687–688, 2013, doi: 10.1126/science.1239276.

[41] F. Jabr, “The connectome debate: Is mapping the mind of a worm worth it,” Scientific American, vol. 18, 2012.

[42] E. Jonas and K. P. Kording, “Could a Neuroscientist Understand a Microprocessor?,” PLoS Comput. Biol., vol. 13, p. e1005268, 2017, doi: 10.1371/journal.pcbi.1005268.

[43] J. Kimble and D. Hirsh, “The postembryonic cell lineages of the hermaphrodite and male gonads in Caenorhabditis elegans,” Dev. Biol., vol. 70, no. 2, pp. 396–417, Jun. 1979, doi: 10.1016/0012-1606(79)90035-6.

[44] T. N. Kipf and M. Welling, “Semi-supervised classification with graph convolutional networks,” arXiv preprint 1609.02907, 2016.

[45] K. P. Kording, G. Blohm, P. Schrater, and K. Kay, “Appreciating the variety of goals in computational neuroscience,” arXiv, [q-bio.NC], 2020.

[46] M. Liu, S. Kumar, A. K. Sharma, and A. M. Leifer, “A high-throughput method to deliver targeted optogenetic stimulation to moving C. elegans populations,” PLoS Biol., vol. 20, e3001524, 2022, doi: 10.1371/journal.pbio.3001524.

[47] W. Lotter, G. Kreiman, and D. Cox, “Deep Predictive Coding Networks for Video Prediction and Unsupervised Learning,” arXiv [cs.LG], 2016, 1605.08104.

[48] H. Markram, “The human brain project,” Sci. Am., vol. 306, pp. 50–55, 2012, doi: 10.1038/scientificamerican0612-50.

[49] B. Mulcahy et al., “Post-embryonic remodeling of the C. elegans motor circuit,” Curr. Biol., vol. 32, pp. 4645–4659.e3, 2022, doi: 10.1016/j.cub.2022.09.065.

[50] A. A. Prinz, D. Bucher, and E. Marder, “Similar network activity from disparate circuit parameters,” Nat. Neurosci., vol. 7, pp. 1345–1352, 2004, doi: 10.1038/nn1352.

[51] F. Rosenblatt, The perceptron, a perceiving and recognizing automaton Project Para, Cornell Aeronautical Laboratory, 1957.

[52] F. Rosenblatt, “The Perceptron: A Probabilistic Model For Information Storage And Organization In The Brain,” Psychological Review, vol. 65, no. 6, pp. 386–408, 1958.

[53] L. Ripoll-Sánchez et al., “The neuropeptidergic connectome of C. elegans,” bioRxiv, 2022.10.30.514396, 2022, doi: 10.1101/2022.10.30.514396.

[54] D. E. Rumelhart, G. E. Hinton, and R. J. Williams, “Learning representations by back-propagating errors,” Nature, vol. 323, no. 6088, pp. 533–536, Oct. 1986.

[55] G. P. Sarma et al., “OpenWorm: overview and recent advances in integrative biological simulation of,” Philos. Trans. R. Soc. Lond. B Biol. Sci., vol. 373, 2018, doi: 10.1098/rstb.2017.0382.

[56] S. Schneider, J. H. Lee, and M. W. Mathis, “Learnable Latent Embeddings for Joint Behavioural and Neural Analysis,” Nature, May 2023, doi: 10.1038/s41586-023-06031-6.

[57] I. H. Stevenson and K. P. Kording, “How advances in neural recording affect data analysis,” Nat. Neurosci., vol. 14, pp. 139–142, 2011, doi: 10.1038/nn.2731.

[58] J. E. Sulston and H. R. Horvitz, “Post-embryonic cell lineages of the nematode, Caenorhabditis elegans,” Dev. Biol., vol. 56, no. 1, pp. 110–156, Mar. 1977, doi: 10.1016/0012-1606(77)90158-0.

[59] J. E. Sulston, E. Schierenberg, J. G. White, and J. N. Thomson, “The embryonic cell lineage of the nematode Caenorhabditis elegans,” Dev. Biol., vol. 100, no. 1, pp. 64–119, Nov. 1983, doi: 10.1016/0012-1606(83)90201-4.

[60] V. Susoy et al., “Natural sensory context drives diverse brain-wide activity during C. elegans mating,” Cell, vol. 184, pp. 5122–5137.e17, 2021, doi: 10.1016/j.cell.2021.08.024.

[61] B. Szigeti et al., “OpenWorm: an open-science approach to modeling Caenorhabditis elegans,” Front. Comput. Neurosci., vol. 8, p. 137, 2014, doi: 10.3389/fncom.2014.00137.

[62] S. Tremblay, C. Testard, J. Inchauspé, and M. Petrides, “Non-necessary neural activity in the primate cortex,” bioRxiv, 2022.09.12.506984, 2022, doi: 10.1101/2022.09.12.506984.

[63] L. R. Varshney, B. L. Chen, E. Paniagua, D. H. Hall, and D. B. Chklovskii, “Structural properties of the Caenorhabditis elegans neuronal network,” PLoS Comput. Biol., vol. 7, e1001066, 2011, doi: 10.1371/journal.pcbi.1001066.

[64] A. Vaswani et al., “Attention is All You Need,” Advances in Neural Information Processing Systems, 2017, [Online]. Available: https://arxiv.org/abs/1706.03762.

[65] A. J. M. Walhout et al., “Protein interaction mapping in C. elegans using proteins involved in vulval development,” Science, vol. 287, no. 5450, pp. 116–122, 2000.

[66] D. M. Walther et al., “Widespread proteome remodeling and aggregation in aging C. elegans,” Cell, vol. 161, no. 4, pp. 919–932, 2015.

[67] J. G. White, E. Southgate, J. N. Thomson, and S. Brenner et al., “The structure of the nervous system of the nematode Caenorhabditis elegans,” Philos Trans R Soc Lond B Biol Sci, vol. 314, no. 1165, pp. 1–340, 1986.

[68] D. Witvliet et al., “Connectomes across development reveal principles of brain maturation,” Nature, vol. 596, pp. 257–261, 2021, doi: 10.1038/s41586-021-03778-8.

